# Temporal epigenomic profiling identifies AHR and GLIS1 as super-enhancer controlled regulators of mesenchymal multipotency

**DOI:** 10.1101/183988

**Authors:** Deborah Gérard, Florian Schmidt, Aurélien Ginolhac, Martine Schmitz, Rashi Halder, Peter Ebert, Marcel H. Schulz, Thomas Sauter, Lasse Sinkkonen

## Abstract

Temporal data on gene expression and context-specific open chromatin states can improve identification of key transcription factors (TFs) and the gene regulatory networks (GRNs) controlling cellular differentiation. However, their integration remains challenging. Here, we delineate a general approach for data-driven and unbiased identification of key TFs and dynamic GRNs, called EPIC-DREM. We generated time-series transcriptomic and epigenomic profiles during differentiation of mouse multipotent bone marrow stromal cells (MSCs) towards adipocytes and osteoblasts. Using our novel approach we constructed time-resolved GRNs for both lineages. To prioritize the identified shared regulators, we mapped dynamic super-enhancers in both lineages and associated them to target genes with correlated expression profiles. We identified aryl hydrocarbon receptor (AHR) and Glis family zinc finger 1 (GLIS1) as mesenchymal key TFs controlled by dynamic MSC-specific super-enhancers that become repressed in both lineages. AHR and GLIS1 control differentiation-induced genes and we propose they function as guardians of mesenchymal multipotency.

## INTRODUCTION

Understanding the gene regulatory interactions underlying cell differentiation and identity has become increasingly important, especially in regenerative medicine. Efficient and specific reprogramming of cells towards desired differentiated cell types relies on understanding of the cell type-specific regulators and their targets (Rackham et al., 2016). Similarly, knowledge of the regulatory wiring in the intermediate stages might allow controlled partial dedifferentiation, and thereby endogenous regeneration, also in mammals (Aguirre et al., 2014).

Great progress has been made in reconstruction of GRNs for various cell types in recent years. While successful, many of the approaches derive their regulatory interactions from existing literature and databases, which may be limiting as the majority of enhancers harboring transcription factor (TF) binding sites are cell type-specific (Forrest et al., 2014). Thus, the regulatory interactions derived from existing databases and literature might be misleading and are likely to miss important interactions that have not been observed in other cell types. Therefore, context-specific expression data have been used to overcome such biases and allow a data-driven network reconstruction (Janky et al., 2014). In addition, other approaches taking advantage of time-series data, such as Dynamic Regulatory Events Miner (DREM) (Schulz et al., 2012), have been developed to allow hierarchical identification of the regulatory interactions. However, while time-series epigenomic data has been used in different studies to derive time point-specific GRNs (Ramirez et al., 2017) (Goode et al., 2016), systematic approaches that integrate the different types of data in an intuitive and automated way are missing.

The central key genes of biological networks under multi-way regulation by many TFs and signaling pathways were recently shown to be enriched for disease genes and are often controlled through so called super-enhancers (SEs), large regulatory regions characterized by broad signals for enhancer marks like H3 lysine 27 acetylation (H3K27ac) (Galhardo et al., 2015) (Hnisz et al., 2013) (Parker et al., 2013) (Siersbaek et al., 2014). Hundreds of SEs can be identified per cell type, many of which are cell type- or lineage-specific and usually control genes that are important for the identity of the given cell type or condition. Thus, SE mapping and SE target identification can facilitate unbiased identification of novel key genes.

An example of lineage specification events with biomedical relevance is the differentiation of multipotent bone marrow stromal progenitor cells (MSCs) towards two mesenchymal cell types: osteoblasts and bone marrow adipocytes. Due to their shared progenitor cells, there is a reciprocal balance in the relationship between osteoblasts and bone marrow adipocytes. Proper osteoblast differentiation and maturation towards osteocytes is important in bone fracture healing and osteoporosis and osteoblast secreted hormones like osteocalcin can influence insulin resistance (N. K. Lee et al., 2007) (Silva & Kousteni, 2012). At the same time bone marrow adipocytes, that occupy as much as 70% of the human bone marrow (Fazeli et al., 2013), are a major source of hormones promoting metabolic health, including insulin sensitivity (Cawthorn et al., 2014). Moreover, increased commitment of the MSCs towards the adipogenic lineage upon obesity and aging was recently shown to inhibit both bone healing and the hematopoietic niche (Ambrosi et al., 2017).

Extensive temporal epigenomic analysis of osteoblastogenesis has been recently reported (H. Wu et al., 2017). However, a parallel investigation of two lineages originating from the same progenitor cells can help to understand both the lineage-specific and the shared regulators important for their (de)differentiation. To identify shared regulators of adipocyte and osteoblast commitment and to delineate a general approach for systematic unbiased identification of key regulators, we performed time-series epigenomic and transcriptomic profiling at 6 different time points over 15 day differentiation of MSCs towards both adipocytes and osteoblasts. We combine segmentation-based TF binding predictions from time point-specific active enhancer data (Schmidt et al., 2017) with probabilistic modeling of temporal gene expression data (Schulz et al., 2012) to derive dynamic GRNs for both lineages. By merging overlapping SEs identified using H3K27ac signal from different time points we obtained dynamic profiles of SE activity across the two differentiations and use these dynamic SEs to prioritize the key regulators identified through the network reconstruction. With this approach, we identified aryl hydrocarbon receptor (AHR) and Glis family zinc finger 1 (GLIS1) as central regulators of multipotent MSCs under dynamic control from SEs that become repressed in the differentiated cells. The repression of these TFs, and in particular of AHR, allows upregulation of many adipocyte- and osteoblast-specific genes, including *Notch* genes, a family of conserved developmental regulators.

## RESULTS

### A subset of differentially expressed genes are shared between adipocyte and osteoblast differentiation

To identify shared regulators of MSC differentiation towards adipocytes and osteoblasts, and to delineate a general approach for a systematic unbiased identification of key regulators, we performed time-series ChIP-seq and RNA-seq profiling at 6 different time points over 15 days of differentiation of mouse ST2 MSCs (Figure 1A). Using ChIP-seq, genome-wide profiles of three different histone modifications, indicating active transcription start sites (TSS) (H3K4me3), active enhancers (H3K27ac), and on-going transcription (H3K36me3) were generated. These data were complemented by corresponding time-series RNA-seq analysis. The successful differentiations were confirmed by induced expression of known lineage-specific marker genes and microscopic inspection of cellular morphology and stainings (Supplementary Figure S1). Interestingly, profiles of the adipogenic marker genes resembled those reported for the yellow adipose tissue (YAT) found in the bone marrow, rather than classic white adipose tissue (WAT) (Scheller et al., 2016), consistent with ST2 cells originating from bone marrow stroma (Figure S1A). Moreover, the expression profiles of *Sp7* and *Runx2* were consistent to those previously reported for mouse osteoblasts (Yoshida et al., 2012).

**Figure 1.**
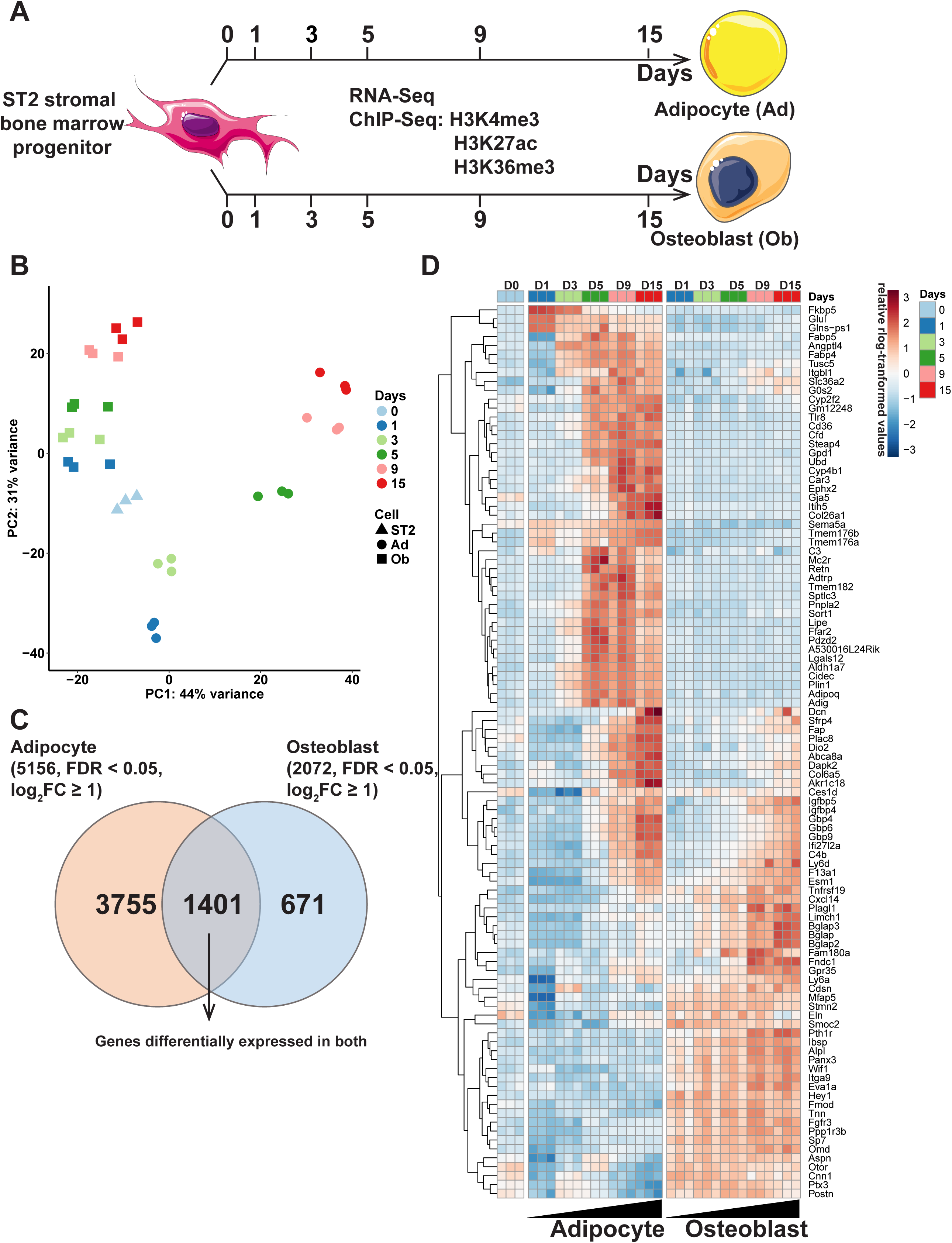
Time-series RNA-seq and ChIP-seq profiling of adipocyte and osteoblast differentiation from shared bone marrow progenitor cells. (A) Schematic representation of the experimental set-up. ST2 MSCs were differentiated towards adipocytes or osteoblasts. Total RNA and chromatin was collected at the indicated time points from both differentiation time courses and subjected to RNA-seq or ChIP-seq analysis using antibodies against the indicated histone modifications.(B) Principle component analysis (PCA) of the RNA-seq data. All replicates are indicated with each time point in a different colour. Triangles marks the undifferentiated ST2 cells, circles the adipocyte (Ad) samples and squares the osteoblast (Ob) samples. (C) Venn diagram comparing all differentially expressed genes at any time point of adipogenesis (log_2_FC > 1, FDR<0.05) to those similarly affected in osteoblastogenesis. (D) Heatmap depicting the top 100 genes with the highest variance across the time points and lineages.

Principal component analysis of the obtained transcriptome profiles confirmed the specification of the cells towards two different lineages with differential temporal dynamics (Figure 1B). Osteoblastogenesis was accompanied by gradual and consistent progression towards a more differentiated cell type while adipogenesis showed more complex dynamics with a big transcriptome shift after one day of differentiation, followed by a more gradual progression during the following days. This is in keeping with the change in the composition of the differentiation medium from day 2 onwards (see Methods for details). In total the adipocyte differentiation was characterized by a total of 5156 significantly differentially expressed genes (log_2_FC≥1, FDR<0.05) across the time series compared to the MSCs (Figure 1C; Supplementary Table S1). During osteoblast differentiation 2072 genes were differentially expressed at least at one time point. 1401 of these genes were affected in both lineages. However, as illustrated by the top 100 genes with highest variance across the time points, which are depicted in the heatmap in Figure 1D, most genes exhibit either lineage-specific or opposing behavior between the lineages. Only a subset of genes showed similar changes in both lineages (Figure 1D). Thus, narrowing the list of genes that could serve as shared regulators of both differentiation or dedifferentiation processes.

### Unbiased data-driven derivation of context-specific dynamic regulatory networks of adipocyte and osteoblast differentiation using EPIC-DREM

In order to take an unbiased and data-driven approach that can benefit from the time series profiles, we have developed a new method to predict condition-specific TF binding using footprint calling in H3K27ac data and TF motif annotation (Gusmao et al., 2016) (Roider et al., 2007). We previously found that using footprints works well for the prediction of gene expression using TEPIC (Schmidt et al., 2017). Our approach uses a randomization strategy, which accurately accounts for differences in footprint lengths and GC-content bias, to assess the significance of TF binding affinity values for each condition or time point (Figure 2A; see Methods for more details). These time point-specific predictions can be combined with the DREM approach (Schulz et al., 2012) to construct lineage-specific networks that are supported by epigenetics data (called EPIC-DREM, Figure 2A).

**Figure 2:**
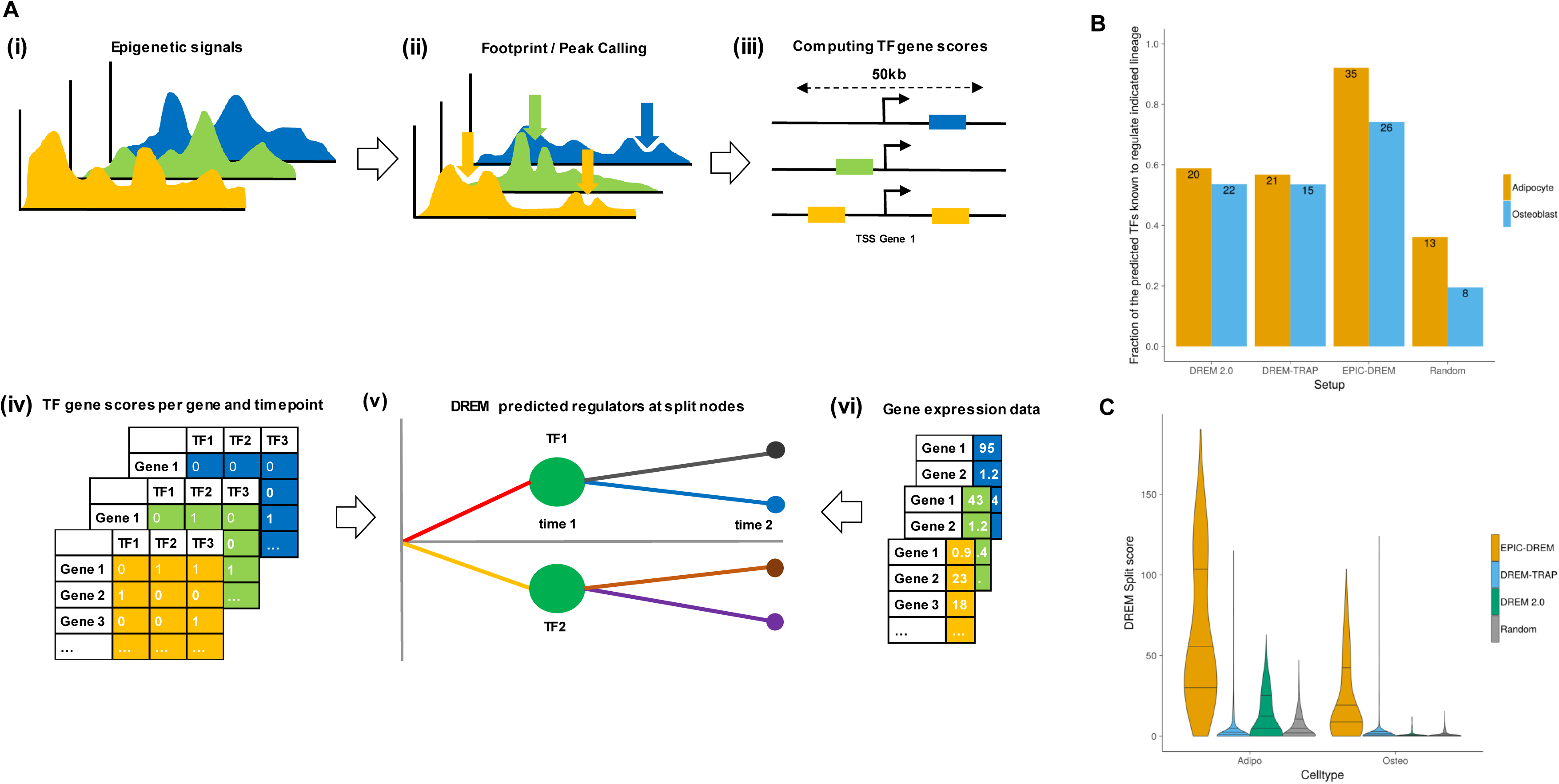
Workflow of the EPIC-DREM approach and benchmarking against other methods. (A) (i-iii) From time-series epigenetic experiments, e.g. DNase1-seq or histone ChIP-seq, putative TF binding sites are identified through footprint/peak calling and are annotated with TF affinities using TRAP. Random genomic regions with similar genomic characteristics (GC-content and length) compared to measured footprints/peaks are generated and annotated with TF affinities as well. A TF- and timepoint-specific affinity threshold can be obtained by applying an empirical *p*-value threshold (e.g. 0.05) on the distribution of TF affinities calculated on the randomly selected regions. Applying the threshold on the TF affinities computed in (iii) results in a set of discrete TF affinities per TF and timepoint. (iv) Using TEPIC, the discrete TF affinities are integrated into discrete, timepoint specific TF-gene association scores. (iv-vi) TF-gene associations and time-series gene expression data are used as input to DREM, which predicts regulators that distinguish subsets of genes that show similar gene expression changes over time. Different time points are indicated by different colours. (B) Literature benchmarking of the different methods. The time-series data from MSC differentation to adipocytes and osteoblasts (Figure 1A) was used as input for 4 different approaches (DREM2.0, EPIC-DREM, DREM-TRAP and random TF-target assignment) to identify top TFs (15 TFs per time point) controlling the respective lineages (see Supplementary Table S2 for the detailed list). For each identified TF the literature was searched for existing evidence for that TFs role in the corresponding differentiation. The fraction of the predicted TFs for which literature exists are shown per method and lineage, with the total number of identified unique top TFs indicated. (C) The distributions of the obtained DREM split scores per method and lineage are shown, with EPIC-DREM typically obtaining highest scores at most split points.

At first we have used 78 TF ChIP-seq datasets from three ENCODE cell lines (GM12878, HepG2, and K562) to test the ability of our approach to prioritize condition-specific TF binding sites using this pipeline. Using a *p*-value cutoff < 0.05 we obtained accurate cell-type specific TF binding predictions with a median precision of ∼70% without a major decrease in recall (Supplementary Figure S2) and, thus, used the same cut-off for our time-series differentiation data.

We used EPIC-DREM to analyse all timepoints of the two differentiation time series of adipogenesis and osteoblastogenesis. Depending on the time point and lineage we predicted 0.6 to 1.4 million footprints per time point, consistent with previous reports on the presence of approximately 1.1 million DNase-seq footprints per cell type (Neph et al., 2012). These footprints were annotated with TFs that are expressed during the differentiation and were associated to target genes within 50 kb of their most 5’TSSs to obtain the TF scores per gene and per time point (see Methods and Figure 2). The full matrix of the time point-specific TF-target gene interactions per lineage can be downloaded at doi:10.5061/dryad.r32t3.

The derived matrix of the predicted time point-specific TF-target gene interactions was combined with the time series gene expression data to serve as input for DREM to identify bifurcation events, where genes split into paths of co-expressed genes (Figures 2 and 3). Knowing the time point-specific TF-target gene interactions allows an accurate association of split points and paths with the key TFs regulating them. To directly test the accuracy and the biological relevance of the EPIC-DREM predictions compared to alternative methods, we performed the same analysis of the time-series expression data also with three alternative approaches for the prediction of TF-gene interactions: 1) based on DREM 2.0, using existing, time-point unspecific ChIP-seq datasets of TF binding sites; 2) DREM-TRAP, where the TF binding site predictions are computed in 2 kb windows centered at the gene TSSs without the time- and context-specific epigenomic profiles, and; 3) Random shuffling of the EPIC-DREM TF-gene interaction matrix (Figures 2B and 2C). To address the biological relevance of these predictions, we considered the top 15 TFs with best split scores from each split at day 0 and combined them into lists of potential master regulators identified for both lineages and each prediction method (Figure 2B, Supplementary Table S2). Next we performed a literature search for each of the predicted TFs on these lists to see whether they have already been associated with adipogenesis or osteoblastogenesis (Supplementary Table S2). From the randomly assigned TFs only 20-30% had been previously found to be associated with the two cell types, while for the TFs predicted by the time-point unspecific methods DREM 2.0 or DREM-TRAP this fraction increased to between 53% and 59%, depending on the lineage (Figure 2B). However, applying EPIC-DREM for the prediction further improved the fraction of biologically relevant known TFs to as high as 92% for adipocyte and 74% for osteoblast differentiation, respectively. Consistently, the DREM enrichment scores obtained by EPIC-DREM were overall higher than those obtained by the other methods (Figure 2C), indicating that the TFs identified by EPIC-DREM can much better explain the observed expression dynamics.

**Figure 3.**
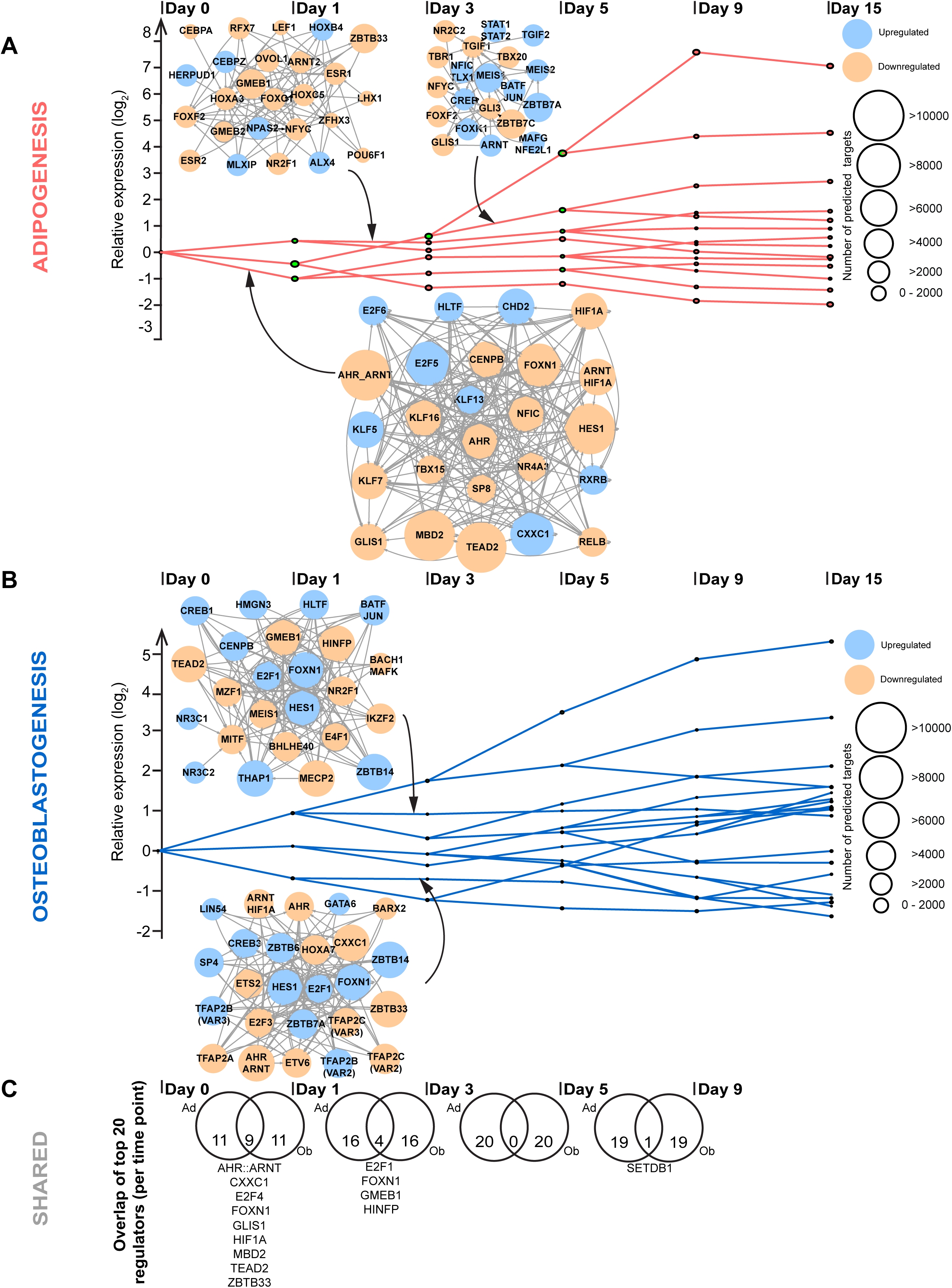
Derivation of lineage-specific regulatory interactions using EPIC-DREM. EPIC-DREM clusters genes into paths of similar expression over time in (A) adipocyte and (B) osteoblast differentiation. Split points differentiate time point-specific changes in gene expression, that are annoted by DREM with matching TFs based on the predicted TF-target gene interactions. For each such path a GRN of thousands of interactions has been derived. The top TFs based on their time point- and path-specific split scores can be used to generate TF-TF networks underlying the observed gene regulation. The TF-TF networks of top 25 TFs are indicated for (A) 3 selected paths in adipocyte and (B) 2 selected paths in osteoblast differentiation. The top TFs are colored depending whether they are upregulated (blue) or downregulated (orange) compared to the previous time point and the size of the node corresponds to the total number of predicted target at the indicated time point. The full set of TFs predicted to control each path with at least 30% of the genes in the upstream split point with a split score ≤ 0.01 are listed in Supplementary Table S3. (C) Overlap of the time point-specific top TFs between the lineages. For both lineages lists of top 20 TFs per time point were generated by combining the predictions from individual splits. The Venn diagrams indicate the extent of overlap between the lineage-specific top TFs at each time point and the shared top TFs are indicated.

DREM clusters co-expressed genes along the time-points and identifies split points with transcriptional regulators assigned to them. Figure 3 shows the split points and the paths of co-expressed genes identified for adipocyte (Figure 3A) and osteoblast (Figure 3B) differentiation. The entire predicted gene regulatory networks consist of up to tens of thousands of nodes and edges, preventing their illustration in an intuitive fashion. However, for the selected DREM paths the TF-TF-networks of the top 25 TFs (based on their split score) are illustrated in Figure 3, with the size of the TF nodes corresponding to the total number of time point-specific predicted target genes across the genome. The full list of TFs controlling the individual paths are provided in Supplementary Table S3 and the full matrix of the gene regulatory networks can be downloaded at doi:10.5061/dryad.r32t3.

Inspection of the TF-TF networks of the identified top TFs confirmed many known positive (e.g. KLF5 on day 0, CEBPA on D1, and TGIF2 on D3) and negative (e.g. HES1 and NR4A3 on day 0, FOXC1 on D1) regulators of adipocyte differentiation (Chao et al., 2008; Farmer, 2006; Horie et al., 2008; Omatsu et al., 2014) (Figure 3A), consistently with the above-mentioned findings that most of the top TFs have been associated with adipogenesis in the literature (Supplementary Table S2). Focusing on the size of the individual nodes in the TF-TF networks directly reveals that TFs controlling the first time point directly after differentiation initiation have the highest number of predicted target genes, with some TFs such as HES1 having more that 10 000 predicted target genes (Figure 3A). Similarly, the results of the regulatory network of osteoblastogenesis confirms several known positive regulators such as the aforementioned HES1, TEAD2, and BHLHE40 (Hakelien et al., 2014; Hassan et al., 2004; Iwata et al., 2006), while revealing many other factors that have not been previously associated to osteoblast differentiation (Figure 3B).

Curiously, EPIC-DREM analysis did not highlight TFs such as PPARG among the top 25 TFs at any bifurcation point, despite PPARG being a well-established master regulator of adipocyte differentiation (Tontonoz & Spiegelman, 2008). To better understand this result and to further explore capabilities of EPIC-DREM we had a detailed look at the predictions related to PPARG and the PPARG:RXRA heterodimer. Indeed, both the PPARG monomer and the heterodimer with RXRA appear among the predicted regulators at most split points (see Supplementary Table S3), in particular for the paths of genes upregulated in adipogenesis. The highest ranking of PPARG is obtained for the genes induced directly after differentiation initiation on day 0, where it is ranked as TF number 34 based on the split score. To better illustrate PPARG regulation in yellow adipocyte differentation, we generated a PPARG-centric GRN on day 3 of differentiation (Supplementary Figure S3), the time point where the highest induction of *Pparg* mRNA takes place (Supplementary Figure S1). Supplementary Figure S3 shows all the TFs predicted to control *Pparg* expression at this time point and the top 200 targets out of the total of 3405 genes predicted to be regulated by PPARG:RXRA (Supplementary Table S4). To see whether these predicted targets are consistent with known direct target genes, we used results from Lefterova *et al*. who experimentally identified PPARG targets as genes with at least 3-fold change in their expression in differentiated adipocytes and with binding of PPARG within 50 kb from the TSS based on ChIP-seq data (Lefterova et al., 2008). Importantly, our predicted PPARG:RXRA targets were significantly enriched for these experimentally validated targets (hypergeometric test p=2.78e-31).

The fact that PPARG is missing from the lists of the very top TFs explaining the gene expression dynamics during adipogenesis could be due to its modest induction during MSC differentiation (Supplementary Figure S1). Indeed, *Pparg* was noted already in earlier work to be weaker induced in bone marrow derived MSCs than in the 3T3-L1 cells that are more commonly used to study white adipocyte differentiation (Cawthorn et al., 2012). In keeping with these observations, analysis of the histone modifications associated with active transcription at the *Pparg* locus in published data from 3T3-L1 cells confirmed the appearance of strong enhancers (based on H3K27ac signal), increased transcription (H3K36me3), and activation of both alternative transcription start sites (H3K4me3) after differentiation initiation (Supplementary Figure S4; (Mikkelsen et al., 2010)). In contrast, in MSCs *Pparg* transcription was abundant already prior to differentiation (H3K36me3), with only one TSS marked by H3K4me3 (transcript variant 1) and strong enhancer regions active across the locus (H3K27ac) (Supplementary Figure S4A). Moreover, while also SE regions could be identified at the *Pparg* locus, consistently with previous work (Hnisz et al., 2013), these did not change significantly during MSC differentiation. Thus, our data confirms PPARG as an important SE-controlled regulator with thousands of putative targets in MSCs and the bone marrow adipocytes, albeit with only modest changes during differentiation.

One of our main aims in analyzing two parallel lineages was the identification of shared regulators that could control the differentiation or dedifferentiation of both cell types. With this aim in mind, we combined the predictions from all bifurcation points per time point to generate separate lists of top 20 TFs with most targets for each time point in both lineages (Figure 3C, see Methods). Next the top 20 TFs at corresponding time points in both lineages were overlapped to uncover the extent and identity of the shared top TFs. As might be expected, the highest level of overlap between the regulators was observed at early differentiation with 9 out of 20 top TFs being same for both lineages (Figure 3C). These are likely to include TFs important for maintaining the multipotent state of MSCs and thus negative regulators of both differentiation processes. Indeed, TFs like the AHR:ARNT heterodimer, E2F4, GLIS1, HIF1A, and TEAD2 have already been associated with gene regulation in different stem cell types (Forristal et al., 2010; Gialitakis et al., 2017; S. Y. Lee et al., 2017; Lv et al., 2017; Maekawa et al., 2011; Singh et al., 2011; Tamm et al., 2011). Interestingly, at day 1 of differentiation the number of shared TFs decreases to four and at later time points the only shared factor is SETDB1, better known as a co-regulator (Frietze et al., 2010). The only TF to appear as shared regulator at two different time points is FOXN1, which is highly connected in GRNs of both lineages, strongly downregulated in adipogenesis but highly expressed in osteoblasts, suggesting FOXN1 might play opposite roles in the two lineages (Figure 3 and Supplementary Table S1).

Taken together, the EPIC-DREM approach can identify many known key regulators of adipocyte and osteoblast differentiation and predicts additional novel regulators and the bifurcation events they control in an entirely unbiased manner relying only on the available time series data.

### Identification of dynamic SEs in adipocyte and osteoblast differentiation

While EPIC-DREM can efficiently identify many of the main regulators of the differentiation time courses, it still yields tens of TFs predicted to control thousands of target genes, most of which are likely to contribute to the cellular phenotype. However, testing tens of different TFs is often not practical. In order to further prioritize the identified main regulators, we hypothesized that the key TFs of the differentiation processes would be under high regulatory load and controlled by SEs with dynamic profiles. To obtain such profiles of SEs across time points, we first identified all SEs with a width of at least 10 kb separately in each of the 10 H3K27ac ChIP-seq data sets from the two time courses. Next, to allow a quantification of the SE signals across time, while also accounting for the changes in the width of the SEs, we combined all SEs from all time points with at least 1 nucleotide overlap into one broader genomic region, called a merged SE. Figure 4A illustrates how the 49 SEs found at the *Cxcl12* locus, producing a chemokine essential for maintenance of the hematopoietic stem cells (HSCs) (Ambrosi et al., 2017), can be combined into one exceptionally large merged SE region that enables the quantification of the SE signal across time. The normalized read counts in the merged SE region were collected and normalized relative to D0 (Figure 4B). To confirm that the obtained profile is a reasonable estimate of the SE activity and to see whether it could indeed be regulating the *Cxcl12* gene expression, we compared the SE signal profiles to the *Cxcl12* mRNA profiles. As shown in Figure 4B, in both lineages the SE signal closely followed the mRNA expression profile with Pearson correlation coefficients of 0.83 and 0.89, respectively. Moreover, similar correlations could be seen for most other identified merged SEs, further confirming the applicability of our approach.

**Figure 4.**
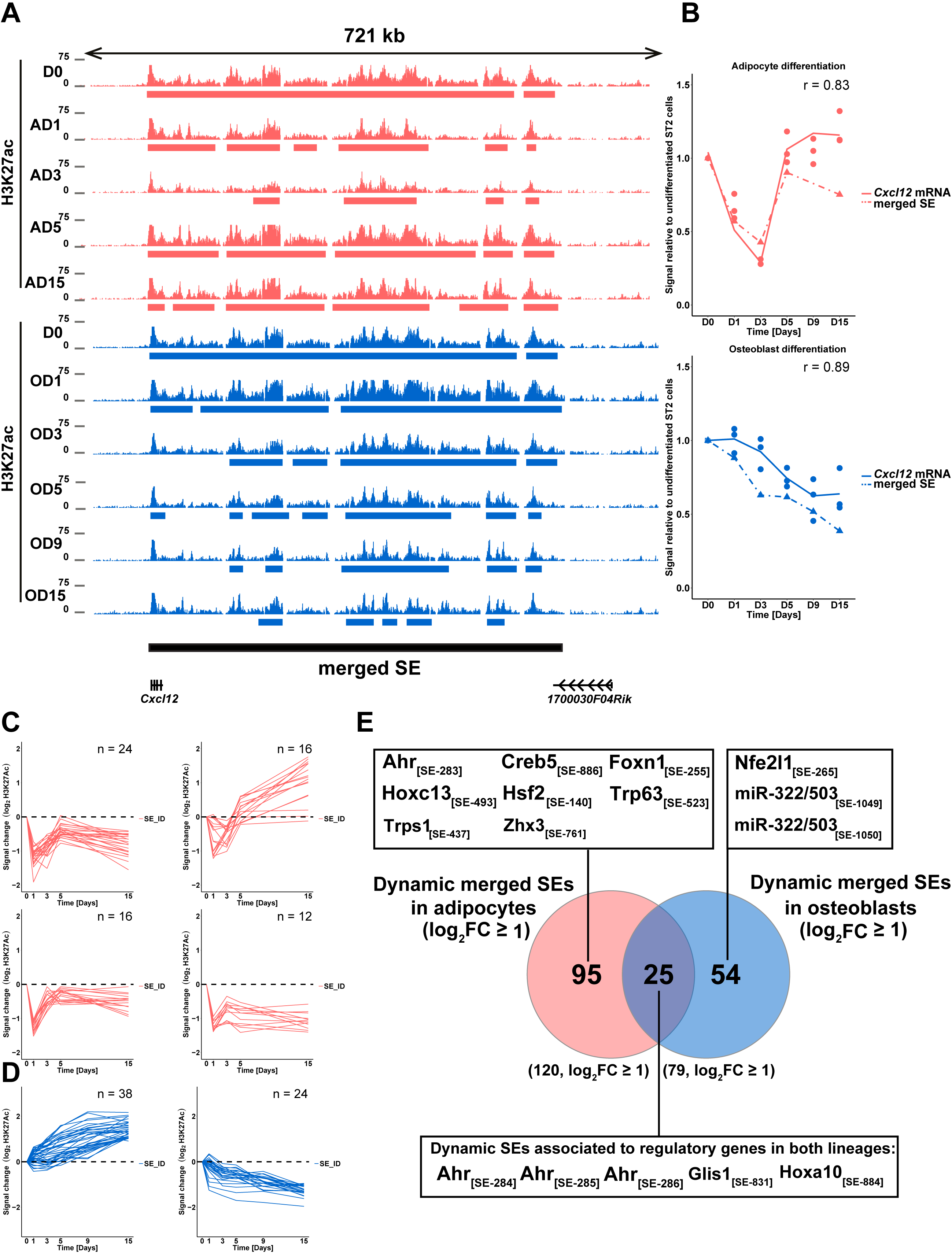
Identification of dynamic SEs. (A) Overview depicting the enrichment of H3K27ac at the *Cxcl12* locus across the time points of adipogenesis (in magenta) and osteoblastogenesis (in light blue). The magenta and light blue bars indicate the SE regions identified by HOMER in adipogenesis and osteoblastogenesis, respectively, and black bar indicates the merged SE derived through overlapping of the individual SEs across the lineages. (B) The correlation between mRNA levels and SE signal. The *Cxcl12* mRNA levels as measured by RNA-seq and the merged SE signal at the *Cxcl12* locus as measured by the reads detected in the H3K27ac IPs are depicted across the time-series of adipocyte (upper panel) and osteoblast (lower panel) differentiation. Intact line = mRNA level and dashed line = merged SE signal. r = Pearson correlation co-efficient (C-D) STEM clustering of the merged SEs according to their dynamic profiles identifies (C) 4 main profiles in adipogenesis and (D) two main profiles in osteoblastogenesis. The additional profiles are shown in Supplementary Figure S5. y-axis indicates the signal change in log_2_-scale. (E) Venn analysis of the dynamic merged SEs from both lineages identifies 95 adipocyte-specific, 54 osteoblast-specific, and 25 shared dynamic SEs with log_2_FC≥1. The dynamic SEs were associated to their target genes based on Pearson correlation with gene expression levels across the time series and the regulatory genes (TFs and miRNAs) associated with dynamic SEs are indicated in the corresponding boxes. All dynamic SEs and their putative target genes are listed in Supplementary Table S6. Adipocyte D9 sample for H3K27ac was not included in the above analysis due to low number of mappable high quality reads.

In total, we identified 1052 merged SEs across the two lineages (Supplementary Table S5). 120 and 79 merged SEs showed a dynamic profile (log_2_FC≥1 in at least one time point) in adipocyte and osteoblast differentiation, respectively. Of these, 25 dynamic SEs showed concordant changes in both differentiation processes (Figure 4C-4E; Supplementary Table S6). Consistent with the different dynamics in transcriptomic changes (Figure 1), the adipocyte SEs could be divided into four separate main profiles based on their dynamics (Figure 4C; Supplementary Figure S5) while most osteoblast SEs could be assigned into one of two simple profiles that either increase or decrease in signal over time (Figure 4D).

In order to identify the genes regulated by these dynamic SEs we calculated the Pearson correlation for all of the dynamic SEs and all of their putative target genes with their TSS located within +/- 500 kb, and associated the SEs to the genes with the highest correlation coefficient (Supplementary Table S6). Out of the 151 genes we found to be associated with the dynamic SEs within the defined distance, 12 were known regulatory genes, classified either as TFs (Heinaniemi et al., 2013) or microRNA genes (Figure 4E). From the 12 regulatory genes, 7 were associated with SEs dynamic only in adipogenesis, namely, *Creb5*, *Hoxc13*, *Hsf2*, *Trp63*, *Trps1*, *Zhx3*, and *Foxn1*. From these FOXN1 was already highlighted among the top TFs by EPIC-DREM analysis, while HOXC13 and HSF2 were also predicted to regulate genes induced during adipogenesis (Supplementary Table S3). The regulatory genes associated with SEs dynamic only in osteoblastogenesis were *Nfe2l1* and *miR-322/503*. From these NFE2L1 was induced and, together with its dimerization partner MAFG, predicted by EPIC-DREM as positive regulator of both osteoblast and adipocyte differentiation (Figure 3, Supplementary Table S3), while miR-322 and miR-503 have been shown to enhance osteogenesis (Gamez et al., 2013; Sun et al., 2017).

Finally, three of the regulatory genes, *aryl hydrocarbon receptor* (*Ahr*), *GLIS family zinc finger 1* (*Glis1*), and *Homeobox A10* (*Hoxa10*), were associated with SEs dynamic in both lineages (Figure 4E). Interestingly, all three TFs are predicted as regulators of MSC differentiation by EPIC-DREM (Supplementary Table S3), and both AHR and GLIS1 belong to the top TFs found to be shared between the two lineages (Figure 3C). AHR, the AHR:ARNT heterodimer, and GLIS1 belong to the TF-TF network controlling genes in early adipogenesis and depicted in Figure 3A. In addition GLIS1 is also included in the TF-TF network controlling genes induced on day 3 of adipogenesis. Similarly, AHR and AHR:ARNT are found in the TF-TF network controlling genes after day 1 of osteoblastogenesis (Figure 3B). Moreover, *Ahr* is associated with four separate merged SEs, more than any other TF in our analysis, and predicted to function upstream of GLIS1 in MSCs (Figure 3A and 4E). Therefore, we next focused on *Ahr* and *Glis1* regulation.

### *Ahr* and *Glis1* are controlled by SEs in MSCs and repressed with lineage-specific dynamics

The four adjacent merged SEs (SE_283_, SE_284_, SE_285_, and SE_286_) at the *Ahr* locus together cover a continuous region over 300 kb of active enhancer signal downstream of the *Ahr* gene in the MSCs (Figure 5A and 5B). SE_831_ covers a region of 29 kb in the 3’end of the *Glis1* gene in the same cells (Figure 6A and 6B). All four SE regions at the *Ahr* locus follow similar dynamics across the time points (Figure 5A-5D) although SE_283_ was identified as being dynamic only in adipogenesis due the FC cut-off in the initial analysis in Figure 4. Moreover, all four SEs showed a very high correlation (r≥0.95) with *Ahr* mRNA levels as measured by RNA-seq (Figure 5C-5D, upper panels) and validated by RT-qPCR (Figure 5C-5D, lower panels). In adipogenesis *Ahr* expression is repressed already by D1 and remains repressed throughout the differentiation, while the signal from all the SE regions is also becoming reduced already after 1 day. In contrast in osteoblasts, the SE regions first show an increase in signal on D1, followed by gradual reduction from D3 onwards, consistent with the *Ahr* mRNA levels.

**Figure 5.**
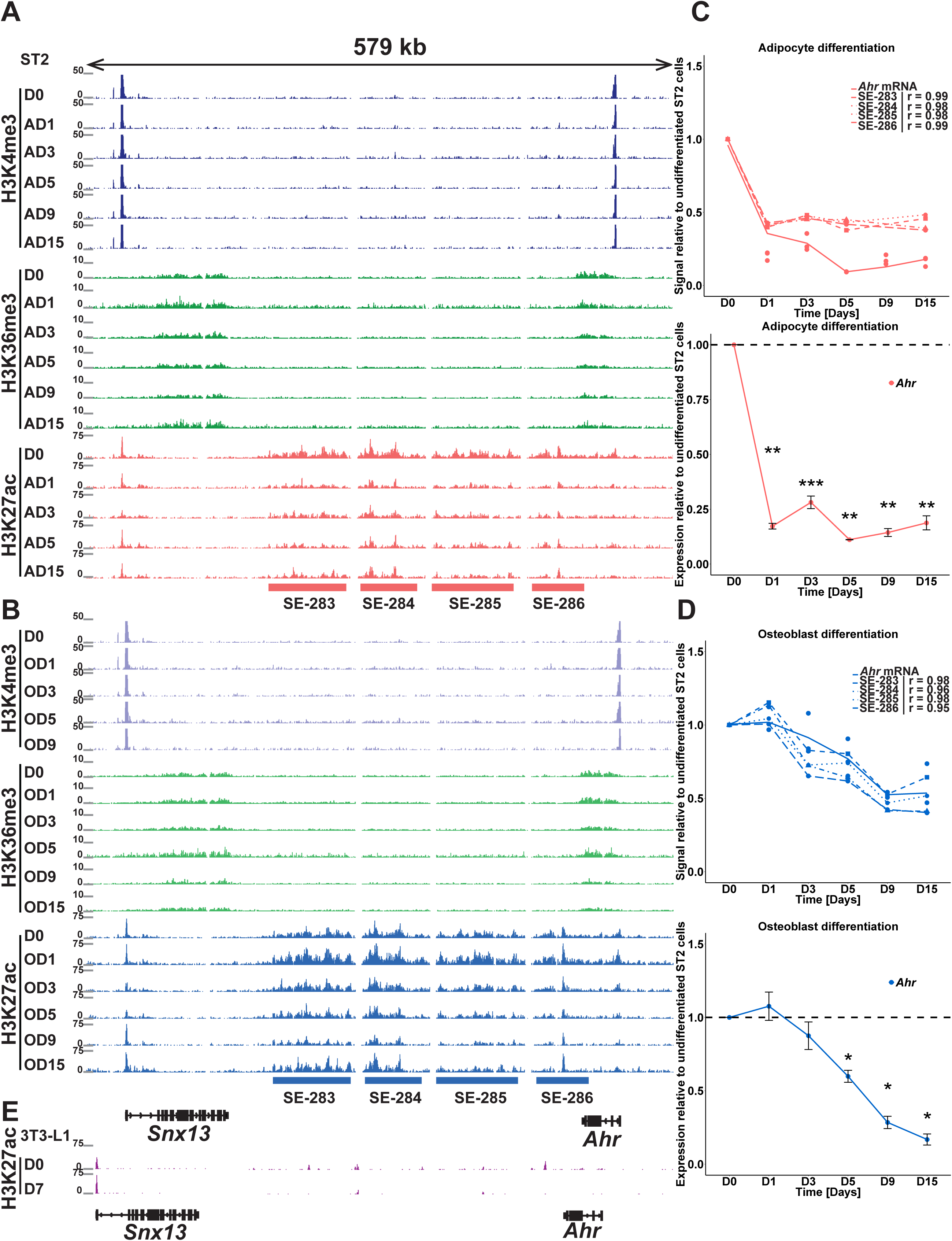
*Ahr* is regulated by multiple SEs with lineage-specific dynamics. (A-B) Overview depicting the enrichments of H3K4me3 (in dark blue and purple), H3K36me3 (in light and dark green), and H3K27ac (magenta and light blue) at the *Ahr* locus across the time points of adipogenesis (A) and osteoblastogenesis (B), respectively. The magenta and light blue bars indicate the merged SE regions identified through the analysis described in Figure 4. See also Supplementary Figure S4. (C-D) *Ahr* downregulation correlates with a decreased signal from all four SEs. The *Ahr* mRNA level was measured across the differentiation by RNA-seq (upper panel) and RT-qPCR (lower panel) in both adipocyte (C) and osteoblast (D) differentiation and is indicated as the intact line. Dashed lines represent the signal from the individual indicated merged SEs. r = Pearson correlation co-efficient. The statistical significance for RT-qPCR measurements compared to the value on D0 was determined by two-tailed Student’s t-test. *=p<0.05, **=p<0.01 and ***=p<0.001. Data points represent mean of 3 biological replicates +/- SEM. AD9 sample for H3K27ac and OD15 for H3K4me3 were not included in the above analysis due to lower number of mappable high quality reads. (E) Overview depicting the enrichment of H3K27ac at the *Ahr* locus in the confluent undifferentiated (D0) and differentiated (D7) 3T3-L1 adipocyte cell line. No SE formation could be detected in these more lineage-committed cells. The data were obtained from (Mikkelsen et al., 2010). The H3K27ac enrichments at the corresponding locus in human cell types are indicated in Supplementary Figure S7A.

**Figure 6.**
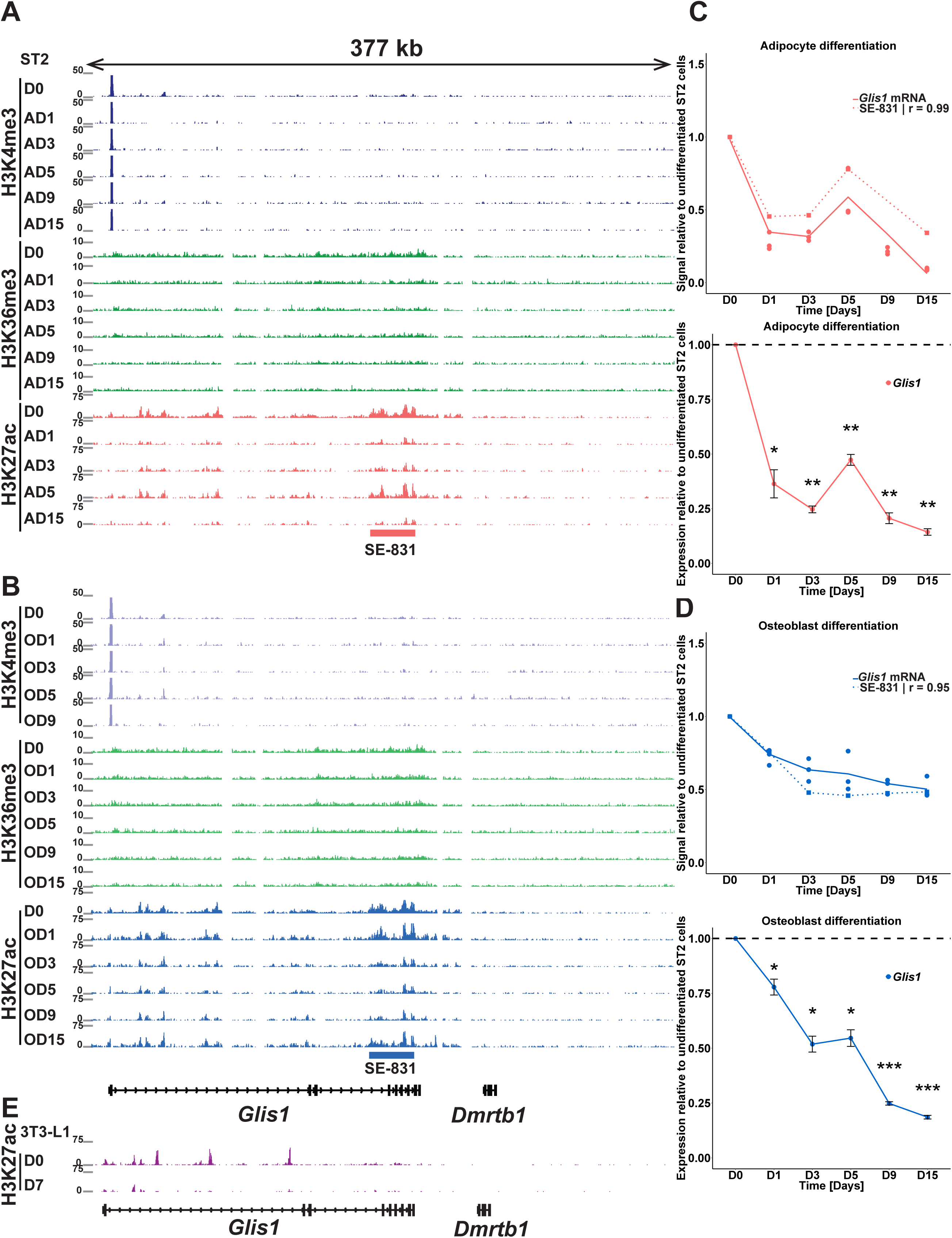
*Glis1* is regulated by a SE with lineage-specific dynamics. (A-B) Overview depicting the enrichments of H3K4me3 (in dark blue and purple), H3K36me3 (in light and dark green), and H3K27ac (magenta and light blue) at the *Glis1* locus across the time points of adipogenesis (A) and osteoblastogenesis (B), respectively. The magenta and light blue bars indicate the merged SE regions identified through the analysis described in Figure 4. See also Supplementary Figure S6. (C-D) *Glis1* downregulation correlates with the signal from SE_831_. The *Glis1* mRNA level was measured across the differentiation by RNA-seq (upper panel) and RT-qPCR (lower panel) in both adipocyte (C) and osteoblast (D) differentiation and is indicated as the intact line. Dashed line represents the signal from the SE_831_. r = Pearson correlation co-efficient. The statistical significance for RT-qPCR measurements compared to the value on day 0 was determined by two-tailed Student’s t-test. *=p<0.05, **=p<0.01 and ***=p<0.001. Data points represent mean of 3 biological replicates +/- SEM. AD9 sample for H3K27ac and OD15 for H3K4me3 were not included in the above analysis due to lower number of mappable high quality reads. (E) Overview depicting the enrichment of H3K27ac at the *Glis1* locus in the confluent undifferentiated (D0) and differentiated (D7) 3T3-L1 adipocyte cell line. No SE formation could be detected in these more lineage-committed cells. The data were obtained from (Mikkelsen et al., 2010). The H3K27ac enrichments at the corresponding locus in human cell types are indicated in Supplementary Figure S7B.

Similarly, the *Glis1* expression changes show high correlation (r≥0.95) with the signal of SE_831_ in both lineages and with both RNA-seq and RT-qPCR (Figure 6C-6D). While in osteoblast differentiation *Glis1* is gradually decreased, in adipocyte differentiation a more dynamic pattern of downregulation is observed with a temporary induction on day 5.

Finally, for both genes and in both lineages the repression is accompanied by a decreased signal of H3K36me3 in the gene bodies of the two genes, confirming their repression at the transcriptional level (Figure 5A-B and 6A-B).

While the newly identified SE clusters at the *Ahr* locus are also flanked by another active gene, *Snx13*, it is unlikely that the SEs contribute to its regulation; during the two differentiations *Snx13* shows very few changes in its expression or H3K36me3 signal (Figure 5A-5B and data not shown). Moreover, inspection of Hi-C data surrounding the *Ahr* locus in different mouse cell types indicates that *Ahr* and the four SEs are located in their own topological domain (TAD) separate from the *Snx13* gene (Supplementary Figure S6) (Tremethick, 2007). Similarly, SE_831_ could be confirmed to be located in the same TAD with *Glis1* and the neighboring *Dmrtb1* gene, which is silenced in MSCs, suggesting that *Glis1* is the main target of the SE.

Interestingly, the SEs controlling *Ahr* and *Glis1* appear to be specific for the multipotent MSCs as only weak or no H3K27ac signal could be detected in the corresponding genomic region in mouse 3T3-L1 pre-adipocytes that are more committed towards the white adipocyte lineage (Figure 5E and 6E, (Mikkelsen et al., 2010)). Importantly, the large SE domains downstream of the *Ahr* gene and the enhancer signals at the 3’end of *Glis1* could be identified also in human MSCs, but not in other inspected human cell types, suggesting that the complex regulation of *Ahr* and *Glis1* expression in these multipotent cells could be conserved and relevant also in human development (Supplementary Figure S7).

### AHR and GLIS1 regulate mesenchymal multipotency through repression of lineage-specific genes

Based on the above results, we reasoned that *Ahr* and *Glis1* could play important roles in maintaining MSCs in a multipotent state. Indeed, previous work has separately shown that both adipogenesis and osteoblastogenesis can be inhibited by toxic compounds like dioxin that are xenobiotic ligands of AHR (Alexander et al., 1998), (Naruse et al., 2002), (Korkalainen et al., 2009), while GLIS1 has been shown to promote reprogramming of fibroblasts to induced pluripotent stem cells (iPSCs) (Maekawa et al., 2011).

To get the first idea whether the two TFs could influence the multipotent state of the MSCs, we performed a knock-down (KD) of AHR and GLIS1 in the MSCs and tested the expression of lineage-specific marker genes used earlier to confirm MSC differentiation (Figure 7A, Supplementary Figure S8A-C and S1). For AHR a KD of approximately 50% could be confirmed at both mRNA and protein level, while Glis1 reduction was around 30% and could not be confirmed at the protein level due to lack of a specific antibody. Still, GLIS1-KD led to a modest but significant induction of *Cebpa*, *Lpl* and *Bglap* expression, while AHR-KD only affected one of the marker genes (*Lpl*). Thus, both AHR and GLIS1 might contribute to maintenance of the MSC state.

**Figure 7.**
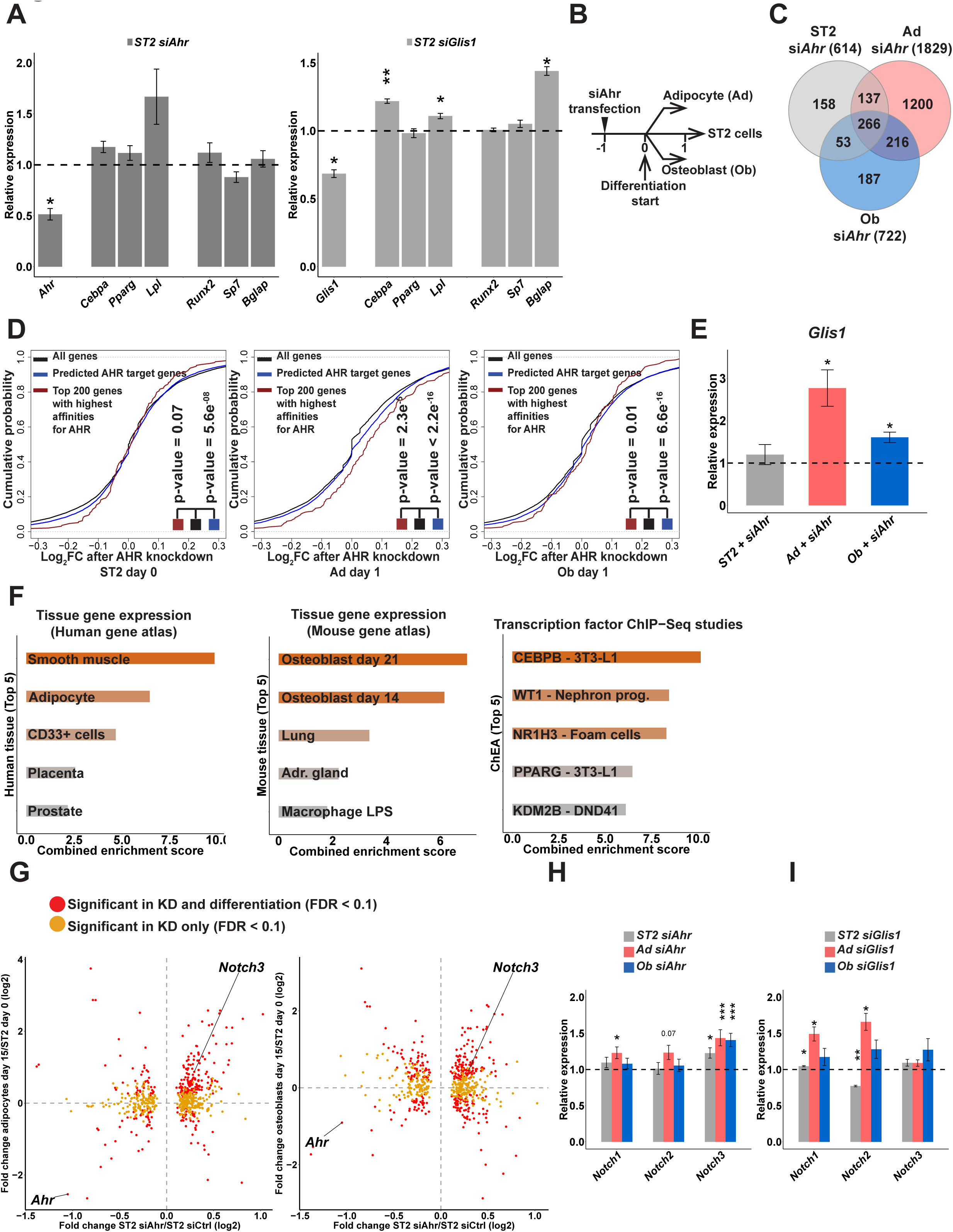
AHR and GLIS1 KDs lead to induction of lineage-specific genes and confirm EPIC-DREM predicted AHR targets. (A) Relative expression levels of differentiation marker genes following AHR or GLIS1 KD in undifferentiated MSCs are shown (y-axis). The statistical significance for RT-qPCR measurements of KD cells against cells transfected with siControl was determined by a two-tailed Student’s t-test (*=p<0.05, **=p<0.01 and ***=p<0.001). Data points represent mean of 3 biological replicates +/- SEM. (B) Schematic representation of the KD RNA-seq experiments. See Supplementary Figure S8 for more details on KD efficiency. (C) Venn diagram comparing the differentially expressed genes from the three different AHR KD conditions (FDR<0.1) identified 266 genes consistently deregulated in all conditions. (D) Confirmation of EPIC-DREM predictions. Cumulative distribution of all expressed genes (black line), all EPIC-DREM predicted AHR target genes at the corresponding condition (blue line), and top 200 targets of AHR (red line) in relation to their log_2_ FC upon AHR depletion per condition as indicated. The significance of the increased FC of the AHR targets was confirmed with the Kolmogorov-Smirnov test and the corresponding p-values are indicated. (E) Relative expression levels of *Glis1* mRNA following AHR KD in the three indicated conditions. See panel A for more details on RT-qPCR. (F) Enrichment analysis of the 266 AHR targets identified in panel C for their preferred tissue expression profiles in human and mouse, and proteins identified as shared regulators in existing ChIP-seq studies. Top 5 most enriched hits are shown for each type of enrichment. X-axis indicates the combined enrichment score (Chen et al., 2013) and coloring the enrichment p-value. (G) Scatter plots indicating the transcriptome-wide log_2_ FCs as measured by RNA-seq in undifferentiated ST2 cells with reduced AHR levels (x-axis) in comparison to the log_2_ FCs of the same transcripts in D15 differentiated adipocytes (left panel) or osteoblasts (right panel). The colour code separates transcripts expressed differentially only in AHR-KD (yellow), or both in KD and in differentiation (red). (H) *Notch3* is induced upon AHR downregulation in all tested conditions, as measured by RNA-seq, while (I) *Notch1* and *Notch2* are affected by GLIS1 KD. See panel A for more details on RT-qPCR. See also Supplementary Figure S8 for expression dynamics of *Notch* receptors during differentation.

While AHR is best known for its role as a xenobiotic receptor, it has been recently suggested that it could also play a role in stem cell maintenance in HSCs under the control of endogenous ligands (Gasiewicz et al., 2014). Moreover, since GLIS1-KD was not more robust and could not yet be confirmed on the protein level, we next focused on the AHR-KD. To directly test whether the endogenous activity of AHR is important for the maintenance of the appropriate transcriptional program of the MSCs, we knocked down AHR in undifferentiated cells, and those differentiated for one day towards either lineage (Figure 7B, Supplementary Figure S8B). Two days following the KD total RNA was extracted and subjected to RNA-seq analysis.

The number of genes affected by the AHR-KD (FDR≤0.1) compared to the respective control siRNA transfections was greatly dependent on the cellular condition and ranged from 614 genes in the undifferentiated cells to 722 and 1819 genes in the one day differentiated osteoblasts and adipocytes, respectively (Figure 7C, Supplementary Table S7). The higher extent of genes affected in the differentiated cells is well in keeping with the number of genes normally changing in the early osteoblasto- or adipogenesis, with adipogenesis associated with more changes (Figure 1). This suggests that early changes induced by the knock-down in the undifferentiated cells are amplified in the differentiating cells.

At first, we took advantage of these context-specific KD data to ask whether the EPIC-DREM predicted primary AHR targets at the corresponding time points were indeed affected by depletion of AHR. As shown in Figure 7D, at each condition the predicted AHR targets were significantly more affected by AHR-KD than all genes on average. Especially the top genes with highest affinity scores for AHR regulation on day 1 of adipogenesis were clearly shifted towards more upregulation at each condition (to the right in the cumulative distribution plot), arguing for functional relevance of the EPIC-DREM predictions and for AHR’s role as transcriptional repressor. As highlighted earlier in Figure 3A, AHR was predicted by EPIC-DREM to act as a direct regulator of *Glis1* in MSCs and early adipogenesis. To test this specific prediction we performed RT-qPCR for *Glis1* in all AHR-KD samples and confirmed a significant change in *Glis1* levels upon AHR depletion in both differentiating adipocytes and osteoblasts (Figure 7E).

To better understand the function of the putative AHR target genes in the MSCs, we overlapped the differentially expressed genes from the undifferentiated cells with those identified in the two other KD experiments and obtained 266 high-confidence target genes that were affected in all three conditions (Figure 7C). Enrichment analysis for tissue-specific expression profiles of these genes in human and mouse gene atlas databases revealed smooth muscle, adipocytes, and osteoblasts as the most enriched cell types, where the AHR regulated genes are normally expressed (Figure 7F). Consistently, inspection of publicly available ChIP-seq data from various cell types revealed the other regulators of the AHR targets to include CEBPB, PPARG, and NR1H3, all of which are important regulators involved in induction of genes during adipocyte differentiation (Galhardo et al., 2014) (Nielsen et al., 2008).

To further elucidate the role of AHR in the undifferentiated cells and the potential impact on lineage-specific genes, we asked how the deregulated genes were expressed in normally differentiated day 15 adipocytes and osteoblasts. The scatter plots in Figure 7G indicate the transcriptome changes taking place upon AHR-KD in MSCs compared to changes of the same genes in the differentiated cell types. As indicated by the color coding, approximately half of the genes affected by AHR-KD were also differentially expressed after normal differentiation (286 in adipocytes and 314 in osteoblasts). Moreover, genes that were upregulated in AHR-KD were preferentially induced in the differentiated cell types (Figure7G, upper right quartile). Therefore, AHR might serve as a guardian of the multipotent state in the undifferentiated cells, with its downregulation allowing increased expression of the lineage-specific genes.

Taken together, the above findings support a role for AHR as a regulator of lineage-specific genes that need to remain repressed in the multipotent MSCs. Among such genes induced upon AHR-KD we identified *Notch3*, a known regulator of cellular differentiation (Bray, 2016) (Figure 7G-H). In both differentiation time courses *Notch3* expression showed an anti-correlated profile compared to *Ahr*, with *Notch3* becoming induced while *Ahr* levels decreased (Figure 5 and Supplementary Figure 8D-E). The *Notch3* induction was also accompanied by increased *Notch4* levels in both lineages while the third abundantly expressed receptor, *Notch1*, was concomitantly downregulated (Supplementary Figure S8D-E). Finally, to see whether the impact of AHR on *Notch3* is mediated via GLIS1, we tested by RT-qPCR the effect of GLIS1-KD on three of the *Notch* genes in the corresponding conditions (Figure 7I). Interestingly, no effect could be seen on *Notch3* expression upon GLIS1-KD while both *Notch1* and *Notch2* were modestly affected. Thus, it appears that a reprogramming of Notch signaling could be involved in the commitment of MSCs and this rewiring is at least partially under AHR- and GLIS1-mediated control.

## DISCUSSION

The ability to obtain unbiased GRNs and to identify their key nodes for any given cell state transition in a data-driven manner is becoming increasingly relevant for regenerative and personalized medicine. Understanding such dynamic networks can be improved by obtaining genome-wide time-series data sets such as transcriptomics or epigenomics data. To seamlessly integrate such data sets we have combined time point-specific high accuracy TF binding predictions with probabilistic modeling of temporal gene expression data and applied it to our own time-series data from mesenchymal differentiation (Figure 2 and Figure 3). Similar time series data collections have previously been used to study for example hematopoiesis (Goode et al., 2016) and myeloid differentiation (Ramirez et al., 2017). However, the derived dynamic GRNs have relied on experimentally identified TF binding sites covering only a fraction of all TF-target gene interactions, or on a sub-network of selected TF-TF interactions, respectively.

EPIC-DREM can reveal the key TFs controlling co-expressed gene sets of interest. Still, consistent with the co-operative nature of TF activity (Siersbaek et al., 2014), the number of putative master regulators is often very large. Recent work elucidating the role of SEs in controlling cell type-specific master regulators has provided researchers with a new tool for data-driven identification of such regulators (Hnisz et al., 2013) (Parker et al., 2013). We hypothesized that merged SEs with dynamic behavior during differentiation based on their H3K27ac signal would allow finding the genes, including the TFs, most relevant for the dynamic process. Indeed, quantification of merged SEs shows high correlation with expression levels of their target genes over time, both validating the approach and allowing for more accurate association of SEs to their target genes (Figure 4). A recent study applied a similar strategy for SE quantification and further showed that SE dynamics, as measured by MED1 occupancy, were predictive of enhancer looping to target genes, and highlighted H3K27ac as the histone modification that best predicted such loop dynamics, further supporting the validity of our approach (Siersbaek et al., 2017).

Combining EPIC-DREM results and dynamic SE profiling points towards several TFs with potentially significant roles in both lineages, including *Foxn1*, *Ahr* and *Glis1* (Figure 4). In addition, dynamic SE profiling supports a role for *Hoxa10*, although in EPIC-DREM analysis it did not make it among the top TFs. Both *Ahr* and *Glis1*, and their SEs, show an overall reduction in signal during both differentiations, although with differential and lineage-specific dynamics (Figure 5 and 6), suggesting that they could play an important role in maintaining MSCs. On the contrary, the SE at the *Foxn1* locus shows a decrease in its signal in adipocyte differentiation (data not shown), while in osteoblast differentiation the signal is further increased. Thus suggesting a role as a positive driver of osteoblast differentiation, while potentially having a negative role in adipocytes. Interestingly, FOXN1 has not been previously shown to be involved in MSC differentiation and our results warrant a further investigation of FOXN1 function in adipocyte and osteoblast differentiation.

Also GLIS1 has not been functionally associated to adipocyte or osteoblast differentiation although recent work has implicated it as differentially expressed in brown adipocyte differentiation (Pradhan et al., 2017). However, consistent with a potential role in the multipotent progenitors, GLIS1 has been shown capable of promoting reprogramming of fibroblasts to induced pluripotent stem cells (iPSCs) (Maekawa et al., 2011). Interestingly, our analysis for enhancer signals at the Glis1 locus in different human cell types confirmed the presence of active enhancers also in human MSCs but not in embryonic stem cells (ESCs) (Supplementary Figure S7). Thus, the ability of GLIS1 to promote cellular reprogramming of iPSCs might reflect its endogenous functions in multipotent stem cells like MSCs, rather than the pluripotent ESCs. Our results implicate GLIS1 as a regulator of MSCs functioning downstream of AHR with its reduction influencing several lineage-specific genes and some of the *Notch* family genes (Figure 7). In addition, both network analysis and the expression dynamics suggest a role for GLIS1 during adipogenesis (Figure 3A and 6). Further work will be needed to elucidate the role of GLIS1 in MSCs and adipocyte differentiation.

Unlike for GLIS1 and FOXN1, previous work has already linked AHR separately to inhibition of both adipocyte and osteoblast differentiation through studies on biological impact of dioxin, an environmental toxin capable of activating AHR (Alexander et al., 1998), (Naruse et al., 2002), (Korkalainen et al., 2009). In 3T3-L1 adipocytes this inhibition is known to be mediated through overexpression of *Ahr* in a dioxin-independent manner (Shimba et al., 2001), while increased levels of *Ahr* expression in MSCs in rheumatoid arthritis are inhibitory of osteogenesis (Tong et al., 2017). Still, the biological function of AHR in the MSCs has remained unclear while maintenance of HSCs, located in the same niche of bone marrow with MSCs, has been suggested to depend on the normal function of AHR (Gasiewicz et al., 2014). Moreover, as HSC maintenance also depends on the cytokines and chemokines provided by the MSCs, AHR is likely to impact HSCs through its gene regulatory functions in both MSCs and HSCs (Ambrosi et al., 2017) (Boitano et al., 2010) (Jensen et al., 2003).

Here we show that repression of *Ahr* in MSC differentiation happens with lineage-specific dynamics and is accompanied by similar reduction in signal of an exceptionally large SE cluster downstream of *Ahr*. Together *Ahr* and the SEs form their own TAD in mouse cells and in human cells the SE signal is specific for MSCs. Analyzing the contributions of the different constituents of the *Ahr*-SEs will be important for understanding which pathways converge to regulate *Ahr* in MSCs and whether they function in a synergistic manner as suggested for some other large SEs (Hnisz et al., 2015) (Dukler et al., 2016).

A detailed characterization and validation of the function of AHR, and other candidate TFs, in MSCs will require the generation of a knock-out system where the TF can be fully depleted in the multipotent cells in an inducible manner. Nevertheless, a reduction of AHR expression in the undifferentiated and early stage differentiated cells already confirmed many of the EPIC-DREM predictions and also revealed an enrichment for lineage-specific genes among AHR targets, including *Notch3* (Figure 6). Notch signaling has been implicated in numerous developmental processes with highly diverse outcomes (Bray, 2016) and also in MSCs *Notch3* regulation is accompanied by changes in other *Notch* genes (Supplementary Figure S8). Interestingly, AHR has been recently linked to regulation of Notch signaling in mouse lymphoid cells and testis (J. S. Lee et al., 2011) (Huang et al., 2016). However, the affected Notch receptors varied depending on the cell type in question. *Notch* genes show different expression profiles across tissues and cell types and *Ahr*-mediated regulation of Notch signaling could be context-specific depending on the prevailing GRN or chromatin landscape. Curiously, *Notch3* has a cell type-selective expression profile, favouring mesenchymal tissues like bone, muscle and adipose tissue (C. Wu et al., 2016).

Our approach for identification of the dynamic GRNs and SEs allows key regulator identification in various time series experiments involving cell state changes. Our current results together with previous data identify AHR and GLIS1 as likely guardians of mesenchymal multipotency, implicate additional TFs as novel regulators of adipocyte and osteoblast differentiation, and provide an extensive resource for further analyses of mesenchymal lineage commitment.

## METHODS

### Cell culture

The mouse MSC line ST2, established from Whitlock-Witte type long-term bone marrow culture of BC8 mice (Ogawa et al., 1988), was used during all experiments. Cells were grown in Roswell Park Memorial Institute (RPMI) 1640 medium (Gibco, Life Technologies, 32404014) supplemented with 10% fetal bovine serum (FBS) (Gibco, Life Technologies, 10270-106, lot #41F8430K) and 1% L-Glutamine (Lonza, BE17-605E) in a constant atmosphere of 37°C and 5 % CO_2_. For differentiation into adipocytes and osteoblasts, ST2 cells were seeded 4 days before differentiation (day-4), reached 100% confluency after 48 hours (day-2) and were further maintained for 48 hours post-confluency (day 0). Adipogenic differentiation was subsequently initiated on day 0 (D0) by adding differentiation medium I consisting of growth medium, 0.5 mM isobutylmethylxanthine (IBMX) (Sigma-Aldrich, I5879), 0.25 µM dexamethasone (DEXA) (Sigma-Aldrich, D4902) and 5 µg/mL insulin (Sigma-Aldrich, I9278). From day 2 (D2) on differentiation medium II consisting of growth medium, 500 nM rosiglitazone (RGZ) (Sigma-Aldrich, R2408) and 5 µg/mL insulin (Sigma-Aldrich, I9278) was added and replaced every 2 days until 15 days of differentiation. Osteoblastic differentiation was induced with growth medium supplemented with 100 ng/mL bone morphogenetic protein-4 (BMP-4) (PeproTech, 315-27). Same media was replaced every 2 days until 15 days of osteoblastogenesis.

### Gene silencing

Undifferentiated ST2 cells (day-1) were transfected with Lipofectamine RNAiMAX (Life Technologies, 13778150) according to manufacturer’s instructions using 50 nM of gene-specific siRNAs against mouse *Ahr* (si*Ahr*) (Dharmacon, M-044066-01-0005), *Glis1* (si*Glis1*) (Dharmacon, M-065576-01-0005) or 50 nM of a negative control siRNA duplexes (si*Control*) (Dharmacon, D-001206-14-05). Cells were collected 48 h post-transfection. Sequences of the siRNAs are listed in Supplementary Table S8.

### Western blotting

After a washing step of the cells with 1x PBS, and addition of 1x Läemmli buffer, the lysates were vortexed and the supernatants were heated at 95°C for 7 minutes. Proteins were subjected to SDS-PAGE (10% gel) and probed with the respective antibodies. The following antibodies were used: anti-AHR (Enzo Life Biosciences, BML-SA210-0025), anti-ACTIN (Merck Millipore, MAB1501). HRP-conjugated secondary antibodies were purchased from Cell Signaling. Signals were detected on a Fusion FX (Vilber Lourmat) imaging platform, using an ECL solution containing 2.5 mM luminol, 100 mM Tris/HCl pH 8.8, 0.2 mM para-coumaric acid, and 2.6 mM hydrogenperoxide.

### RNA extraction and cDNA synthesis

Total RNA was extracted from ST2 cells using TRIsure (Bioline, BIO-38033). Medium was aspirated and 1000 µL of TRIsure was added to 6-wells. To separate RNA from DNA and proteins, 200 µL of chloroform (Carl Roth, 6340.1) was added. To precipitate RNA from the aqueous phase, 400 µL of 100% isopropanol (Carl Roth, 6752.4) was added and RNA was incubated at −20°C overnight. cDNA synthesis was done using 1 µg of total RNA, 0.5 mM dNTPs (ThermoFisher Scientific, R0181), 2.5 µM oligo dT-primer (Eurofins MWG GmbH, Germany), 1 U/µL Ribolock RNase inhibitor (ThermoFisher Scientific, EO0381) and 1 U/µL M-MulV Reverse transcriptase (ThermoFisher Scientific, EP0352) for 1h at 37°C or 5 U/ µL RevertAid Reverse transcriptase for 1 h at 42°C. The PCR reaction was stopped by incubating samples at 70°C for 10 minutes.

### Quantitative PCR

Real-time quantitative PCR (qPCR) was performed in an Applied Biosystems 7500 Fast Real-Time PCR System and using Thermo Scientific Absolute Blue qPCR SYBR Green Low ROX Mix (ThermoFisher Scientific, AB4322B). In each reaction 5 µL of cDNA, 5 µL of primer pairs (2 µM) and 10 µL of the Absolute Blue qPCR mix were used. The PCR reactions were carried out at the following conditions: 95°C for 15 minutes followed by 40 cycles of 95°C for 15 seconds, 55°C for 15 seconds and 72°C for 30 seconds. To calculate the gene expression level the 2^-(ΔΔCt)^ method were used where ΔΔCt is equal to (ΔCt_(target gene)_ – ΔCt_(housekeeping gene)_)_tested condition_ - (ΔCt_(target gene)_ – ΔCt_(housekeeping gene)_)_control condition_. *Rpl13a* was used as a stable housekeeping gene and D0 or si*Control* were used as control condition. Sequences of the primer pairs are listed in Supplementary Table S8.

### Chromatin Immunoprecipitation

Chromatin immunoprecipitation of histone modifications was performed on indicated time points of adipocyte and osteoblast differentiation. Cells were grown on 10 cm^2^ dishes. First, chromatin was cross-linked with formaldehyde (Sigma-Aldrich, F8775-25ML) at a final concentration of 1% in the culture media for 8 minutes at room temperature. Then, the cross-linked reaction was quenched with glycine (Carl Roth, 3908.3) at a final concentration of 125 mM for 5 minutes at room temperature. The formaldehyde-glycine solution was removed and cells were washed twice with ice-cold phosphate-buffered saline (PBS) (Lonza, BE17-516F) containing cOmplete^TM^ mini Protease Inhibitor (PI) Cocktail (Roche, 11846145001). Then, cells were lysed in 1.7 mL of ice-cold lysis buffer [5 mM 1,4-Piperazinediethanesulfonic acid (PIPES) pH 8.0 (Carl Roth, 9156.3); 85 mM potassium chloride (KCl) (PanReac AppliChem, A2939); 0.5 % 4-Nonylphenyl-polyethylene glycol (NP-40) (Fluka Biochemika, 74385)] containing PI and incubated for 30 minutes on ice. The cell lysates were then centrifuged at 660 xg for 10 min at 7°C and the pellet was resuspended in 400 µL of ice-cold shearing buffer [50 mM Tris Base pH 8.1 (Carl Roth, 4855.2); 10 mM Ethylenediamine tetraacetic acid (EDTA) (Carl Roth, CN06.3); 0.1 % Sodium Dodecylsulfate (SDS) (PanReac Applichem, A7249); 0.5 % Sodium deoxycholate (Fluka Biochemika, 30970)] containing PI. Chromatin was sheared with a sonicator (Bioruptor®Standard Diagenode, UCD-200TM-EX) during 20 cycles at high intensity (30 s off and 30 s on) for the ST2 cells differentiated into adipocytes and osteoblasts and 25 cycles at high intensity (30 s off and 30 s on) for the ST2 differentiated into osteoblasts for 9 days on. The sheared cell lysate was then centrifuged at 20817 xg for 10 minutes at 7°C and the supernatant containing the sheared chromatin was transferred to a new tube. For each immunoprecipitation 10 µg (for H3K4me3) or 15 µg (for H3K27ac and H3K36me3) of sheared chromatin and 4 µg as input were used. The sheared chromatin was diluted 1:10 with modified RIPA buffer [140 mM NaCl (Carl Roth, 3957.2); 10 mM Tris pH 7.5 (Carl Roth, 4855.2); 1 mM EDTA (Carl Roth, CN06.3); 0.5 mM ethylene glycol-bis(β-amino-ethyl ether)-N,N,N’,N’-tetraacetic acid (EGTA) (Carl Roth, 3054.3); 1 % Triton X-100 (Carl Roth, 3051.2); 0.01 % SDS (PanReac Applichem, A7249); 0.1 % sodium deoxycholate (Fluka Biochemika, 30970)] containing PI. The diluted sheared chromatin was incubated overnight with the recommended amount provided by the manufacturer of an antibody against H3K4me3 (Millipore, 17-614), 5 µg of an antibody against H3K27ac (Abcam, ab4729) or 5 µg of an antibody against H3K36me3 (Abcam, ab9050). The next day, the antibodies were captured using 25 µL of PureProteome™ Protein A Magnetic (PAM) Bead System (Millipore, LSKMAGA10) for 2 hours at 4°C on a rotating wheel. After, the PAM beads were captured using a DynaMag^TM^-2 magnetic stand (Life Technologies, 12321D). The supernatant was discarded and the PAM beads were washed twice with 800 µL of Immunoprecipitation wash buffer 1 (IPWB1) [20 mM Tris, pH 8.1 (Carl Roth, 4855.2); 50 mM NaCl (Carl Roth, 3957.2); 2 mM EDTA (Carl Roth, CN06.3); 1 % Triton X-100 (Carl Roth, 3051.2); 0.1 % SDS (PanReac Applichem, A7249)], once with 800 µL of Immunoprecipitation wash buffer 2 (IPWB2) [10 mM Tris, pH 8.1 (Carl Roth, 4855.2); 150 mM NaCl (Carl Roth, 3957.2); 1 mM EDTA (Carl Roth, CN06.3), 1 % NP-40 (Fluka Biochemika, 74385), 1 % sodium deoxycholate (Fluka Biochemika, 30970), 250 mM of lithium chloride (LiCl) (Carl Roth, 3739.1)], and twice with 800 µL of Tris-EDTA (TE) buffer [10 mM Tris, pH 8.1 (Carl Roth, 4855.2); 1 mM EDTA (Carl Roth, CN06.3), pH 8.0]. Finally, the PAM beads and the inputs were incubated with 100 µL of ChIP elution buffer [0.1 M sodium bicarbonate (NaHCO_3_) (Sigma-Aldrich, S5761); 1 % SDS (PanReac Applichem, A7249)]. The cross-linking was reversed by adding 10 µg of RNase A (ThermoFisher, EN0531) and 20 µg of proteinase K (ThermoFisher, EO0491) at 65°C overnight. Then, the eluted chromatin was purified using a MinElute Reaction Cleanup Kit (Qiagen, 28206) according to the manufacturer’s instructions. The DNA concentration was measured using the Qubit® dsDNA HS Assay Kit (ThermoFisher, Q32851) and the Qubit 1.0 fluorometer (Invitrogen, Q32857) according to the manufacturer’s instructions.

### ChIP-Seq

The sequencing of the ChIP samples was done at the Genomics Core Facility in EMBL Heidelberg, Germany. For sequencing, single-end and unstranded reads were used and the samples were processed in an Illumina CBot and sequenced in an Illumina HiSeq 2000 machine. In total, 979 572 918 raw reads were obtained. Raw reads quality was assessed by fastqc [v0.11, (Ruvkun & Hobert, 1998)]. This quality control unveiled that some reads were containing part of the adapters. Those spurious sequences were cleaned up from the genuine mouse sequences by AdapterRemoval (Lindgreen, 2012) [v1.5]. The PALEOMIX pipeline (Schubert et al., 2014) [v1.0.1] was used for all steps from FASTQ files to BAM files including trimming, mapping, and duplicate marking. This workflow ensures that all files are complete and valid. Retained reads were required to have a minimum length of 25 bp. Bases with unreliable Phred scores (0-2) were trimmed out. In total 31 909 435 reads were discarded (3.26%). Eventually, 947 663 483 reads were retained (96.74%). Trimmed reads were further mapped using BWA (Li & Durbin, 2009) [v0.7.10] with the backtrack algorithm dedicated to short sequences. The mouse reference was the mouse genome GRCm38.p3 (mm10, patch 3) downloaded from NCBI. For validating, merging BAM files, and marking duplicates, we used the suite tool Picard [v1.119, (Adams et al., 2000)]. Duplicates were marked but not removed. Only reads with a mapping quality of 30 were retained to ensure a unique location on the genome resulting in 661 364 143 reads (69.79% of the trimmed reads). The samples with a coverage of less than 8 million reads (mapping quality > 30) were excluded from the downstream analysis. Raw FASTQ and BAM files have been deposited in the European Nucleotide Archive with the accession number PRJEB20933.

The ChIP-Seq peaks were called with Model-based analysis of ChIP-Seq (Zhang et al., 2008) (MACS) version 2.1.0 for H3K4me3, with HOMER (Heinz et al., 2010) for H3K27ac, and with SICER (Zang et al., 2009) version 1.1 for H3K36me3, using input from undifferentiated ST2 cells as control for IPs from D0 cells and input from D5 adipocyte- or osteoblast-differentiated cells for the IPs from the respectively differentiated cells.

### RNA-Seq

The sequencing of the time course samples was done at the Genomics Core Facility in EMBL Heidelberg, Germany. For sequencing, single-end and unstranded reads were used and the samples were processed in an Illumina CBot and sequenced in an Illumina NextSeq machine.

The sequencing of the AHR knock-down samples was performed at the Luxembourg Center for Systems Biomedicine (LCSB) Sequencing Facility. The TruSeq Stranded mRNA Library Prep kit (Illumina) was used to prepare the library for sequencing with 1 µg of RNA as starting material according to manufacturer’s protocol. The library quality was checked using an Agilent 2100 Bioanalyzer and quantified using Qubit dsDNA HS assay Kit. The libraries were then adjusted to 4 nM and sequenced on a NextSeq 500 (Illumina) according to the manufacturer’s instructions.

The obtained reads were quality checked using FastQC version 0.11.3 (Ruvkun & Hobert, 1998). Cutadapt version 1.8.1 (Martin, 2011) was used to trim low quality reads (`-q 30` parameter), remove Illumina adapters (`-a` parameter), remove reads shorter than 20 bases (`-m 20` parameter) with an error tolerance of 10% (`-e 0.1` parameter). Then, removal of reads mapping to rRNA species was performed using SortMeRNA (Kopylova et al., 2012) with the parameters `--other`, `--log`, `-a`, `-v`, `--fastx` enabled. Lastly, the reads were quality checked using FastQC version 0.11.3 to control whether bias could have been introduced after the removal of Illumina adapters, low quality reads and rRNA reads. Then, the reads were mapped to the mouse genome mm10 (GRCm38.p3) and using the gene annotation downloaded from Ensembl (release 79) using the Spliced Transcripts Alignment to a Reference (Dobin et al., 2013) (STAR) version 2.5.2b using the previously described parameters (Baruzzo et al., 2017). The reads were counted using the function *featureCounts* from the R package *Rsubread* (Liao et al., 2014) version 1.4.6-p3 and the statistical analysis was performed using DESeq2 (Love et al., 2014) version 1.14.1 in R 3.3.2 and RStudio (RStudio Team (2015). RStudio: Integrated Development for R. RStudio, Inc., Boston, MA).

### EPIC-DREM analysis

To identify TFs that have a regulatory function over time, we designed a new computational workflow that combines the computational TF prediction method TEPIC (Schmidt et al., 2017) with DREM (Schulz et al., 2012), a tool to analyze the dynamics of transcriptional regulation.

We identified TF footprints in the H3K27ac signal using HINT-BC (Gusmao et al., 2016), which is included in the Regulatory Genomics Toolbox, version 0.9.9. Next, we predicted TF binding in those footprints using TEPIC, version 2.0. We used the provided set of 687 PWMs for *Mus musculus* and mouse genome version mm10 (GRCm38) to predict TF affinities using TRAP (Roider et al. 2007) within TEPIC. As DREM requires a time point-specific prediction of binding of a regulator with its target, we needed to develop an approach to determine a suitable TF-specific affinity cut-off, for each time point. For this, we created a similar set of random regions that mirrors the GC content and length distribution of the original sequences of the footprints. TF affinities *a_r_* calculated in the random regions are used to determine a suitable cut-off for the original affinities *a_o_* using the frequency distribution of the TF affinities. Affinities for TF *i* are denoted by *a_ri_* and. *a_o_* Let *r ∈ R* denote a randomly chosen genomic region that is screened for TF binding, and let |*r*| denote its length. Analogously, let *o ∈ O* denote a footprint that is screened for TF binding, and let |*o*| denote its length. We normalize both *a_ri_* and *a_oi_* by the length of their corresponding region and obtain the normalized TF affinities 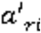 and 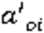:

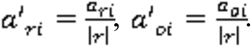

Using the distribution of 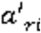 values we derive a TF-specific affinity threshold *t_i_* for a p-value cut-off of *0.05* (See section *ChIP-seq validation of TEPIC affinity cut-off* for how this p-value was chosen). For a TF *i* we compute a binary affinity value *b_oi_* from the original affinity 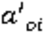 according to the cut-off *t_i_* with:

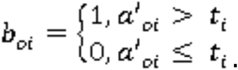

The binary affinity values *b_oi_* can be used to compute a binary TF – gene association *a_gi_* between gene *g* and TF *i*: 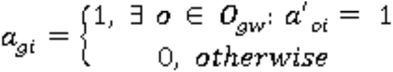, where *o_gw_* denotes all footprint regions that occur within a window of size *w* around the TSS of gene *g*.

Informally, a gene *g* is associated to TF *i* if there is a predicted binding site within a window of predefined size *w* around the gene’s TSS. Here, we use *w = 50kb*.

Together with gene expression estimates, the TF– gene associations can be directly used as input to DREM. In this analysis, we used version 2.0.3 of DREM. The entire workflow of EPIC-DREM is shown in Figure 2A. TEPIC is available online at https://github.com/SchulzLab/TEPIC, DREM can be downloaded at http://www.sb.cs.cmu.edu/drem/.

### ChIP-seq validation of TEPIC affinity cut-off

To validate that the affinity threshold described above indeed results in an adequate separation between bound and unbound sites, we conducted a comparison to TF-ChIP-seq data. We obtained TF-ChIP-seq data from ENCODE for K562 (18 TFs), HepG2 (36 TFs), and GM12878 (24 TFs). In addition, we downloaded H3K27ac data for the mentioned cell lines from ENCODE. A list of all ENCODE accession numbers is provided in Supplementary Table S9. As described above, we called footprints using HINT-BC and calculated TF affinities in the footprints as well as in the randomly selected regions that map the characteristics of the footprints. To understand the influence of different thresholds, we calculated affinity thresholds for the following p-values: 0.01, 0.025, 0.05, 0.075, 0.1, 0.2, 0.3, 0.4, and 0.5. All affinities below the selected affinity value are set to zero, the remaining values are set to one. The quality of the discretization is assessed through the following “peak centric” validation scheme, as used before in (Cuellar-Partida et al., 2012). The positive set of the gold standard is comprised of all ChIP-seq peaks that contain a motif predicted by FIMO (Grant et al., 2011), the negative set contains all remaining ChIP-seq peaks. A prediction is counting as a true positive (TP) if it overlaps the positive set, it counts as a false positive (FP) it if overlaps the negative set. The number of false negatives (FN) is the number of all entries in the positive set that are not overlapped by any prediction. For all TFs in all cell lines we calculate Precision (PR) and Recall (REC) according to

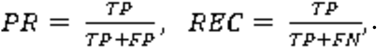

As one can see, Precision is increasing with a stricter p-value threshold, while Recall is decreasing. We found that using 0.05 seems to be a reasonable compromise between Precision and Recall. The median Precision and Recall values calculated over all cell lines and all TFs are shown in Supplementary Figure S2A. Detailed results on a selection of TFs that are present in all three cell lines are shown in Supplementary Figure S2B.

### Generating TF-TF interaction networks for DREM splits

We devised a general strategy to create TF-TF interaction networks for individual DREM splits. For a split of interest, we retrieve the top 25 regulators *T*, ranked according to the DREM split score. For each regulator 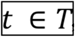, we determine the target genes 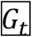 by declaring each gene *g* to be a target of TF *t*, if and only if 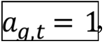, where 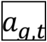 is the binary value for the TF affinity of TF *t* in the vicinity of gene *g*, as introduced above. Here, we want to show only interactions among the top regulators, thus, in order to include a directed edge from *t* to *g* in the TF-TF network, it must hold that. 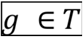 For reasons of simplicity, we limit the number of shown interactions per TF *t* to 10, ranked by the numerical affinity values across all 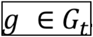.

Additionally, we scale the size of each node representing a regulator 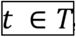, according to its total number of target genes |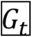|in six discrete levels (<2000, <4000, <6000, <8000, <10 000, and >10 000). This allows an interpretation of the importance of the regulator outside the scope of the interactions among the top TFs for the shown split. The node colour indicates the expression change of the depicted TF compared to time-point zero. Orange refers to downregulation, whereas blue refers to upregulation. The networks are arranged with *graphviz* and the *neato* layout algorithm (Gansner et al., 2005).

### DREM-TRAP and Random TF-gene assignment

To assess the impact of epigenetics data on DREM predictions, we designed a baseline approach, assuming that most transcriptional regulation for a gene is occurring in its promoter region. Hence, for each gene *g*, we consider a 2 kb window around the most 5’ TSS of *g*. Within this window, we compute TF affinities as described above for 687 PWMs for *Mus musculus* and mouse genome version mm10 (GRCm38). Also, we applied the same thresholding approach as described for EPIC-DREM. Thereby, DREM-TRAP represents approaches that are purely sequence and annotation based and do not account for changes in the chromatin.

As another sanity check for EPIC-DREM, we permuted the EPIC-DREM input matrix, which is based on time-point specific footprint calls. In detail, we randomly shuffled column 1 (TF), column 2 (target gene), and column 4 (time point). Thus the number of TFs, target genes, and timepoint entries is not affected, only the mapping between them changed.

### Enrichment score of DREM splits and its aggregation

DREM uses a hypergeometric distribution to assess the association of a TF to genes in a distinct path, the so called split-score, where a lower value means a stronger association. Because we carry out a considerable number of tests per split (as we test more than 600 TFs), we correct the split-scores for multiple testing using Bonferroni correction.

To compare the enrichment scores across the different versions of DREM used in this manuscript, we –log2 transformed the Bonferroni corrected split-scores and show violin plots of the transformed scores, which we call *DREM split score*. For those scores, the higher the value, the better is the association of the TFs identified at split points to genes on the adjacent paths.

To identify essential TFs across multiple splits at one time point, we perform p-value aggregation using Fisher’s method on the split scores, i.e. we compute

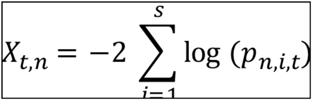

Here, 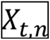 is the aggregated score for TF *t* at time point *n*, *i* is the current split, *s* indicates the number of splits at Timepoint *n* and 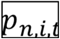 is the split score for TF *t* at split *i* at time point *n.* Next, we rank the 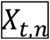 by time point n and thereby obtain a list of top regulators per time point.

### Identification of dynamic merged SEs

In order to identify temporal SEs across both lineages, BedTools (Quinlan & Hall, 2010) version 2.24.0, Hypergeometric Optimization of Motif EnRichment (Heinz et al., 2010) (HOMER) version 4.7.2 and Short Time-series Expression Miner (Ernst & Bar-Joseph, 2006) (STEM) version 1.3.8 were used. First, the coverage of individual SEs was summarized using `genomeCoverageBed` command using `–g mm10` and `–bg` parameters. Then, `unionBedGraphs` command was used to combine multiple SEs coverage into a single map such that the SEs’ coverage is directly comparable across multiple samples. Finally, `mergeBed` command was used to combine SEs overlapping ≥ 1 bp into a single merged SE which spans all the combined SEs. In order to calculate the normalized read count number of merged SEs, `annotatePeaks.pl` with `–size given` and `–noann` parameters was used. Lastly, STEM was used to cluster and identify SEs temporal profiles and SEs with *Maximum_Unit_Change_in_Model_Profiles_between_Time_Points* 2 and *Minimum Absolute Expression Change* 1.0 were considered as dynamic.

For the dynamic SEs we calculated the Pearson correlation for the putative target genes with their TSS located within +/- 500 kb, and associated the SEs to the genes with the highest correlation coefficient.

### Enrichment analysis

EnrichR (14.4.2017) (Chen et al., 2013; Kuleshov et al., 2016) was used to perform gene enrichment analysis.

### Availability of data and materials

The datasets generated and analysed during the current study are available in the European Nucleotide Archive with the accession number PRJEB20933. All scripts and generated GRNs can be retrieved from https://github.com/sysbiolux and from https://github.com/SchulzLab/TEPIC.

## ACKNOWLEDGEMENTS

We would like to thank Dr. Maria Bouvy-Liivrand for help with establishing the ST2 cell culture and differentiation and EMBL Gene Core at Heidelberg for support with high-throughput sequencing. The experiments presented in this paper were carried out using the HPC facilities of the University of Luxembourg (Varrette et al., 2014).

## AUTHOR CONTRIBUTIONS

DG, TS and LS conceived the project and designed the experiments and analysis. DG performed all the experiments and DG and LS analyzed the results. DG and AG performed the RNA-seq and ChIP-seq analysis. FS and MHS developed the EPIC-DREM approach. DG, FS and MHS performed the EPIC-DREM analysis. MS and DG performed the Western blotting. RH prepared the libraries and performed the sequencing for the AHR-KD experiments. PE developed the randomization method to derive control footprint regions. All authors commented on the manuscript.

